# Holistic optimization of an RNA-seq workflow for multi-threaded environments

**DOI:** 10.1101/345819

**Authors:** Ling-Hong Hung, Wes Lloyd, Radhika Agumbe Sridhar, Saranya Devi Athmalingam Ravishankar, Yuguang Xiong, Eric Sobie, Ka Yee Yeung

## Abstract

**Summary:** For many next-generation sequencing pipelines, the most computationally intensive step is the alignment of reads to a reference sequence. As a result, alignment software such as the Burrows-Wheeler Aligner (BWA) is optimized for speed and and is often executed in parallel on the cloud. However, there are other less demanding steps that can also be optimized and significantly increase the speed especially when using many threads. We demonstrate this using a Unique-molecular-identifier (UMI) RNA sequencing pipeline consisting of 3 steps: split, align, and merge. Optimization of all three steps yields a 40% increase in speed when executed using a single thread. However, when executed using 16 threads, we observe a 4-fold improvement over the original parallel implementation and more than an 8-fold improvement over the original single-threaded implementation. In contrast, optimizing only the alignment step results in just a 13% improvement over the original parallel workflow using 16 threads.

## 1. Introduction

Due to advances in next generation sequencing (NGS) there are now more than half a million datasets in the Gene Expression Omnibus (GEO) (Barrett *et al.*, 2012). A major bottleneck for the analyses of these data is aligning the reads to a reference genome. Many efficient methods for sequence alignment or pseudo-alignment have been developed, such as the Burrows-Wheeler Aligner (BWA) (Li and Durbin, 2009), STAR (Dobin *et al.*, 2013), Kallisto (Bray *et al.*, 2016), Salmon (Patro *et al.*, 2017)). With the ready availability of cheap multi-threaded and distributed computing on cloud platforms, the alignment step is often run in parallel, greatly reducing the time required for analyses of the data. However, as noted by Amdahl more than 50 years ago (Amdahl, 1967), there are diminishing returns with greater numbers of threads as the non-parallelizable components eventually become rate-limiting. Optimization and partial parallelization of these less computationally intensive components can yield significant improvement in a highly parallel environment. We demonstrate this with a pipeline for the analyses of Unique Molecular Identifier (UMI) RNA-seq data. In UMI RNA-seq, a sequence tag with a barcode and random sequence identifies which well on the 96 or 384 well plate the read originates from and controls for amplification artifacts (Islam *et al.*, 2013).

## 2 A three-step UMI RNA-seq workflow

The RNA-seq alignment workflow is designed for the Unique Molecular Identifier (UMI) RNA-seq data generated by the LINCS Drug Toxicity Signature (DToxS) Generation Center at Icahn School of Medicine at Mount Sinai in New York (Xiong *et al.*, 2017). The workflow described in the the Standard Operating Procedure (SOP 3.1) and the scripts and supporting files for the analytical workflow originate from the Broad Institute (Soumillon *et al.*, 2014). There are 3 steps in the original pipeline implemented by two Python scripts. The first step (split) takes the sequence tag in the first read and appends it to the sequence identifier in the second read creating a new set of fastq files. The second read contains the actual cDNA sequence derived from the native transcripts. The second step (align) aligns the second reads to the human reference genome using BWA. The third step (merge) takes the resulting SAM files, filters out the counts contributed by reads tagged with identical UMIs and then consolidates the transcript counts for each of the wells. Our optimized pipeline consists of the following three major changes.

1. **Demultiplexing the reads by wells:** In addition to appending the sequence tag to the title of the read, reads from the same wells are combined, resulting in 96 new fastq files. Since each well is an independent experiment, the subsequent steps can operate on these files in parallel. The smaller files also greatly reduced the RAM needed, an important consideration as multi-threaded applications often require more RAM.
2. **Parallelism is increased.**

- **Split:** The original split step was not multi-threaded. The split step now operates on different fastq files simultaneously when there were multiple threads available.
- **Align:** The original align step used BWA aln to generate initial alignments which are piped to BWA samse to combine the results and generate a SAM file. BWA aln can use multiple threads but samse is single-threaded. The new align step spawns multiple instances of BWA, each operating on a different file. This parallelizes both BWA aln and BWA samse.
- **Merge:** The original merge step compiled the counts using a single thread and a single large hash table. The new merge can have threads working simultaneously on different files.
3. **Python scripts are replaced by C++ executables** Even though Python uses the same libraries for CPU-intensive operations such as decompressing files, there is a significant amount of text manipulation that is handled by the Python scripts. Converting to C++ removes the overhead from dynamic typing and automatic garbage collection when using Python.

Additional details of our optimizations are found in the Appendix.

## 3 Benchmarking results

We compared the execution time and memory usage between the optimized and original workflows when the number of threads is varied using an Amazon Web Services (AWS) EC2 spot instance. In Figure 1, we see that when using a single thread the dominant contribution to the execution time is the align step. The situation changes when using 16 threads. The CPU-intensive align step now takes the least amount of time in the original workflow. Improving the parallelization of the align step results in a 70% improvement in this step but only a 13% improvement in the overall workflow. See Table 3 in the Appendix for details. However, optimizing all the steps results in almost a 4-fold improvement in speed over the original parallel workflow and more than an 8-fold increase in speed over the original single-threaded workflow. While memory requirements increase with the number of threads used, this is offset by the memory saved from examining reads from individual wells. The optimized workflow takes less than half the memory of the original workflow even with 16 threads.

## 4 Conclusions

The ready availability of on-demand cloud computing means that computationally demanding workflows are now typically run with multiple threads. Optimization efforts have largely and correctly focused on the most computationally intensive components of the pipeline. However, due to diminishing returns, optimization of other less obvious computational modules for improvement can yield dramatic benefits in a multi-threaded cloud environment.

**Figure 1:**
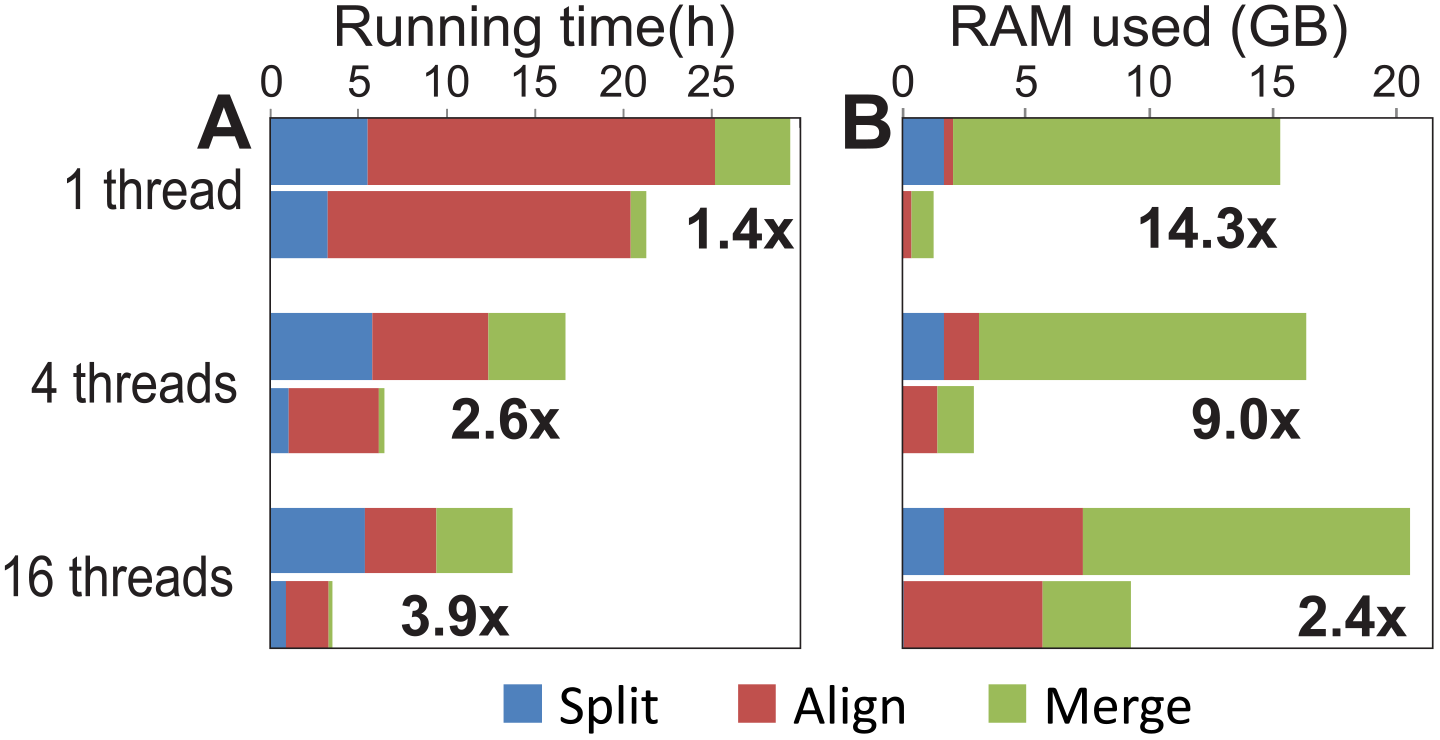
Comparison of execution time (panel A) and memory usage (panel B) between the optimized and original workflows. The upper bars represent the original workflow and the lower bars the optimized workflow. The speedups for 1 thread, 4 threads and 16 threads are shown. All workflows were executed on m4.4xlarge Amazon Web Services (AWS) EC2 spot instances with 64 GB of RAM and 16 vCPU cores. The input data were 6 pairs of fastq files totaling 47 GB on an attached EBS volume. The execution time and memory usage represent the median values across 3 runs. Values are comparisons of the total execution time and the RAM required. The numerical values are in Tables 1 and 2 in Appendix 2

## Acknowledgements

LHH, WL, ES and KYY are supported by NIH grant R01GM126019. LHH and KYY are also supported by NIH grant U54HL127624 and the AMEDD Advanced Medical Technology Initiative. YX and ES are also supported by NIH grant U54HG008098. AWS Cloud Credits for Research (WL and KYY) were used to run the benchmarks.

## Appendix 1: Details of optimized implementation

## 4.1 Original implementation structure

The original implementation consisted of a master bash script which basically handled the setup of the directories and passing of parameters to the two Python scripts that performed the UMI based quantitation of the RNA-seq data. The output files were text files that had the counts for each gene. For the purposes of the benchmarking we moved the system call to BWA from within the Python script to a the shell script. This allowed us to time and measure the memory consumption of that component separately.

## 4.3 Optimized implementation

The optimized workflow duplicated the bash script and replaced the Python scripts with compiled C++ executables that took the same arguments (and have some additional configuration options such as the number of threads).

The details of the implementation for the three components of the workflow, split, align, merge follow:

- **Split:** The split executable reads and decompresses if necessary, a list of pair-end fastq files and splits them into separate fastq files with the sequence tag as part of the title line. Files are organized by the well as specified by the barcode in the sequence tag. The software asks the user for a file which lists the barcodes and wells that they map to. It also asks the user the maximum number of base pair variation due to sequence errors that is tolerated when matching the wells. Since the barcodes are small (6 bases), the split software maps the sequence to an integer and generates a lookup table for all possible tags and maps them to a well when the identity of the well is unambiguous and the variation in base pairs is within tolerance. All other sequences are mapped an “unmatched” well value. This is a constant time operation and is very fast due to the small size of the table. The fiear read sequence is then read and the barcode matched. This is appended to the title line for the read. The actual sequence of from the cDNA is read from the second read. The well is determined from the barcode and the newly constructed read is written to the corresponding fastq file. Unmatched barcodes are written to a unmatched fastq file. This task is parallelized using OpenMP. Each thread operates on a separate pair of input pair-end fastq files and generates a separate set of output well-specific files. The major optimizations are in the parallelization at the level of input files, the fast lookup and resolution of barcodes and the separation of the reads by wells. The last step is a key optimization for later steps.
- **Align:** The align step is performed using a shell script. The script asks the user how many threads to use (parameter nThreads). The master thread assembles a list of all the fastq files from the split step (typically the number of pairs of input files times the number of wells). It then spawns nThread processes that call BWA. Each thread goes through the list of fastq files, and checks whether another thread has written a lock file indicating that another thread is working on that file. When it finds a fastq file to work on, it writes a new lock file to prevent another thread from working on the same file. The added parallelism increases the speed of the align step by 70% when using 16 threads.
- **Merge:** The original merge step did not have reads organized by file and compiled the counts using a single thread and a single large hash table. The new merge uses OpenMP to generate and manage threads that work simultaneously on different files. The division of files by wells also reduces the size of the the hash (since the barcode part of the sequence tag is the same in each file) and the hash table (the number of reads is also reduced). The code further increases the efficiency of hashing by first mapping the sequence tag to an integer and using an unordered_set to check for uniqueness. The original implementation used the sequence concatenated with metadata to form a string and which is hashed default Python text hashing function. Hashing of a long string requires several passes of the hash function and is slower than hashing a 64 bit integer. However, the major savings in speed and memory are due to the division of the data into files sorted by wells. This is especially important when 384 well plates are used instead of 96 well plates and when using many threads, which will consume more memory. A major implementation difference in the merge procedure is the filtering by UMI. The original UMI paper assumes that any additional reads with the same UMI that map to the same position are amplification artifacts and should be discarded. The merge module follows this recipe by default but also allows the user flexibility to exclude reads with the same UMI that map nearby, which could happen, for example, due minor sequencing errors or ambiguities. The original merge implementation excluded duplicate UMI’s that mapped to the same gene.

## 4.4 Testing of optimized implementation

Testing of the optimized implementation is complicated by 3 factors. The first is the difference in the manner that UMI filtering is performed. The second is the fact that BWA will produce slightly different alignments even when the same reads are organized in different files. This is due to the way that it handles the equally scoring alignments by choosing one at random. Finally, the implementations of the sorting by Python and C++ differ, again on how ties are handled.

During initial development, we split the files and filtered duplicate UMI’s in the same manner as the original. This produced the identical counts as the original. Some results were sorted slightly differently due the differences in sorting between Python and C++. However, the splitting and the filtering are key to the optimization and correctness of the new implementation and will produce slightly different results from the original implementation. Consequently, after these features were added, we only tested whether the optimized software produced the same results as previous implementations. For all benchmarks, the output files were compared to the single thread runs and were identical except for the order of items which had not been sorted and were written to the file in a different order by the different threads.

## 4.5 Software availability

Code, supporting scripts and Dockerfiles are available at https://github.com/BioDepot/LINCS_RNAseq_cpp and Docker images at https://hub.docker.com/r/biodepot/rnaseq-umi-cpp/

## Appendix 2: Benchmark data

See Table 1 and Table 2 in the Appendix show the raw median data in Figure 1.

**Table 1:** Execution times in hours:minutes:seconds. The median execution time

| Implementation | Threads | Split time | Align time | Merge time | Total time |
| --- | --- | --- | --- | --- | --- |
| Original | 1 | 5:31:06 | 19:38:09 | 4:16:12 | 29:25:27 |
| Optimized | 1 | 3:14:51 | 17:10:16 | 0:50:37 | 21:15:44 |
| Original | 4 | 5:48:43 | 6:33:31 | 4:19:17 | 16:41:31 |
| Optimized | 4 | 1:04:18 | 5:06:22 | 0:17:59 | 6:28:39 |
| Original | 16 | 5:22:31 | 4:02:41 | 4:17:49 | 13:43:01 |
| Optimized | 16 | 0:53:02 | 2:24:58 | 0:12:43 | 3:30:43 |

**Table 2:**
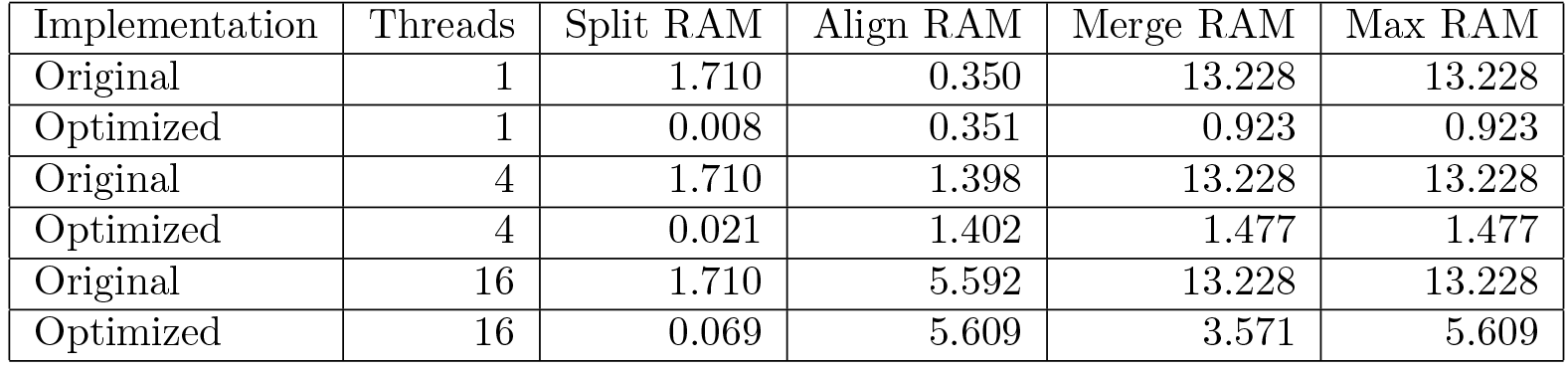
Memory usage in Gigabytes. The median memory usage across three trials are shown.

| Implementation | Threads | Split RAM | Align RAM | Merge RAM | Max RAM |
| --- | --- | --- | --- | --- | --- |
| Original | 1 | 1.710 | 0.350 | 13.228 | 13.228 |
| Optimized | 1 | 0.008 | 0.351 | 0.923 | 0.923 |
| Original | 4 | 1.710 | 1.398 | 13.228 | 13.228 |
| Optimized | 4 | 0.021 | 1.402 | 1.477 | 1.477 |
| Original | 16 | 1.710 | 5.592 | 13.228 | 13.228 |
| Optimized | 16 | 0.069 | 5.609 | 3.571 | 5.609 |

**Table 3:** Effect of optimized computational module (combinations of split, align, merge) on total median run time.

| Optimization | Total run time (16 Threads) |
| --- | --- |
| None | 13:43:01 |
| Split | 9:13:32 |
| Align | 12:05:18 |
| Merge | 9:37:55 |
| Split and Align | 7:35:49 |
| Split and Merge | 5:08:26 |
| Align and Merge | 8:00:12 |
| Split, Align and Merge | 3:30:43 |

